# Lamin regulates the dietary restriction response via the mTOR pathway in Caenorhabditis *elegans*

**DOI:** 10.1101/2020.10.27.357913

**Authors:** Chayki Charar, Sally Metsuyanim-Cohen, Yosef Gruenbaum, Daniel Z Bar

## Abstract

Animals subjected to dietary restriction (DR) have reduced body size, low fecundity, slower development, lower fat content and longer life span. We identified lamin as a regulator of multiple dietary restriction phenotypes. Downregulation of *lmn-1*, the single *Caenorhabditis elegans* lamin gene, increased animal size and fat content, specifically in DR animals. The LMN*-*1 protein acts in the mTOR pathway, upstream to RAPTOR and S6K, key component and target of mTOR complex 1 (mTORC1), respectively. DR excludes the mTORC1 activator RAGC-1 from the nucleus. Downregulation of *lmn-1* restores RAGC-1 to the nucleus, a necessary step for the activation of the mTOR pathway. These findings further link lamin to metabolic regulation.

## Introduction

Dietary restriction (DR) is a metabolic intervention with a conserved response among many organisms. It is one of the most effective methods to prolong lifespan as well as health span in many animal models, with beneficial effects in humans. It reduces many age related pathologies including diabetes and cardiovascular diseases (Colman et al., 2009; Mair and Dillin, 2008).

The mechanistic target of rapamycin (mTOR) is a nutrient sensor that functions as a central regulator of metabolism and physiology. Inhibition of mTOR prolongs the lifespan and improves the healthspan of many model organisms (Johnson et al., 2013; Mannick et al., 2014). Suppression of mTOR is one of the underpinning mechanisms of the beneficiary effects of DR (Papadopoli et al., 2019). By contrast, mTOR signaling is dysregulated in cells harboring disease causing mutations in human lamin A (*LMNA*) genes (Chiarini et al., 2019).

Lamins are type V nuclear intermediate filaments, conserved in metazoan evolution, and key components of the nuclear lamina. Mutations in the human *LMNA* gene cause numerous diseases, including metabolic diseases, accelerated ageing disorders and muscle diseases. *Caenorhabditis elegans*, a free-living nematode with a single lamin gene (*lmn-1*), is extensively used in the research of lifespan regulating pathways. In addition, it is used in the research of the structure and function of the nuclear lamina (Bank et al., 2011; Link et al., 2018; Turgay et al., 2017; Wiesel et al., 2008).

We identified lamin as the regulator of multiple DR phenotypes. Knockdown of *lmn-1* increased fat content and animal size in DR animals, while simultaneous knock-down of RAPTOR (a key component of mTORC1) abolished this size increase. Furthermore, *lmn-1* knockdown had no impact on size in DR animals lacking S6K, one of the main targets of the mTOR. Finally, downregulation of *lmn-1* enabled the nuclear entry of RAGC-1 (Ras related GTP binding C), an essential mechanism for activation of mTORC1 signaling.

## Results

### *lmn-1* regulates size in DR animals

*C. elegans* feeding depends on rhythmic contractions (pumping) of the pharynx (Avery and Horvitz, 1990). *eat-2* (*ad1116*) mutated *C. elegans* (henceforth *eat-2*) have a reduced pumping rate, resulting in slower food uptake. This results in longer life span, smaller body length, lower fat content, and smaller brood size (Bar et al., 2016; Walker et al., 2005), thus serving as a suitable model for DR research (Walker et al., 2005). To gain insight to the roles of lamin in ageing and metabolism, young *eat-2* adults were fed with bacteria expressing *lmn-1* double stranded RNA (RNAi), that effectively knocked-down LMN-1 (**Supp. Fig. 1**). Surprisingly, downregulation of *lmn-*1 significantly increased the body size (length and width) of *eat-2* animals, but not of controls (**Fig. 1*A*** and ***B***, *p*<0.0001). Since the number of somatic cells in *C. elegans* is fixed, *eat-2* animals are smaller due to smaller cell size (Mörck and Pilon, 2007; Walker et al., 2005). To validate that lamin knockdown rescues cell size, we measured the length of cells expressing red fluorescent protein (RFP) fused to the MYO-2 muscle specific protein. Downregulated for *lmn-1* increase in cell length in *eat-2* animals (**Fig. 1*C*** and ***D***, *p*<0.0001). This increase was proportional to the increase seen in the whole entire animal (**Fig. 1*A*** and ***B***). These results suggest that cell size changes constitute the increase in animal size. Of note, downregulation of *lmn-1* did not increase the food intake, as the pumping rate remained unaffected (**Fig. 1*E***).

**Figure 1.**
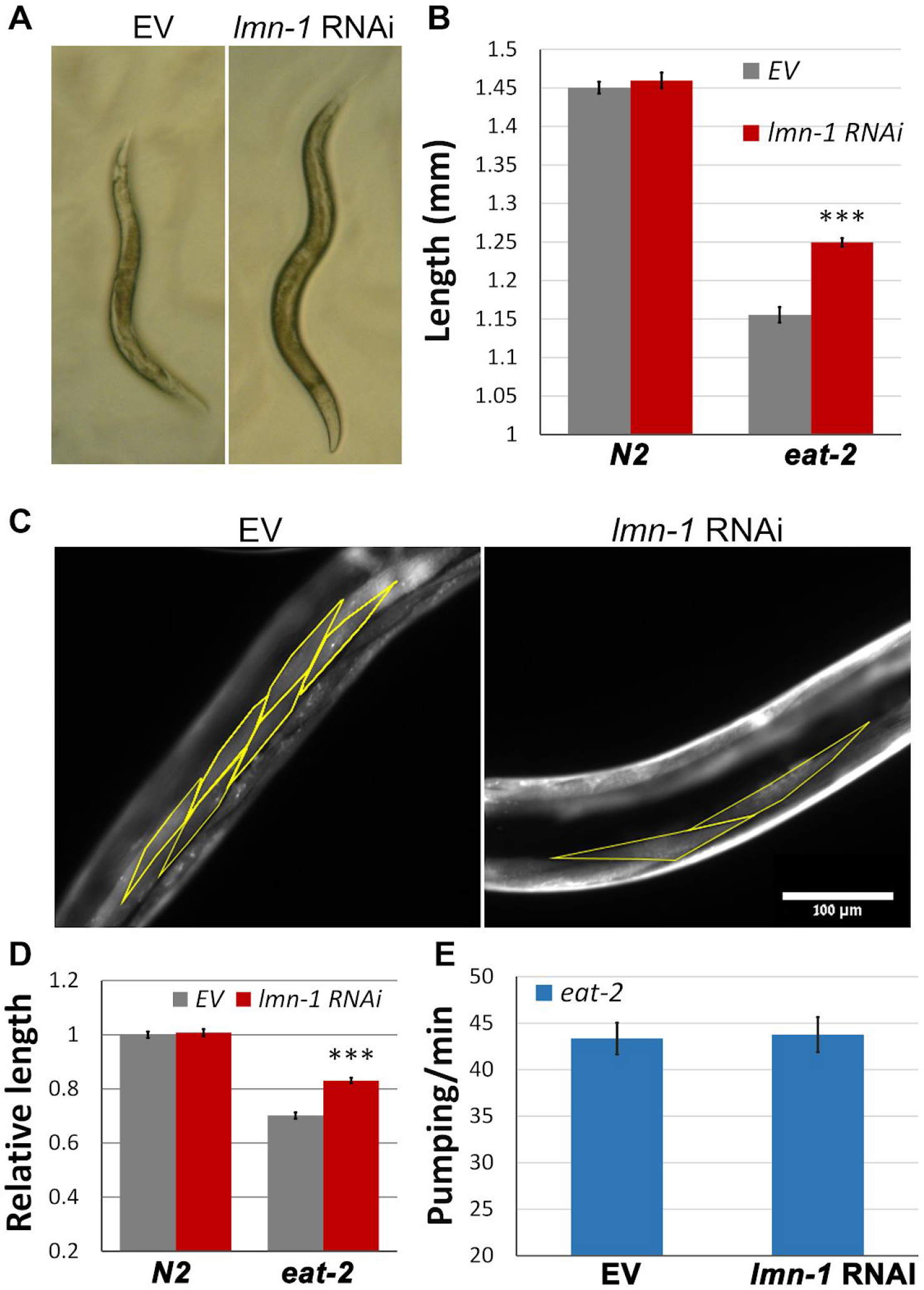
Size reduction in DR worms requires *lmn-1* activity. (*A*) Stereomicroscope images showing *eat-2* (*ad1116*) worms fed for 72h with *lmn-1* (RNAi) or EV. (*B*) Average length of animals from (A). *n*(N2) = 93 and *n*(*eat*-2) = 89 animals. (C) Representative microscope images of *eat-2* (*ad1116*) worms expressing *myo-2*;RFP. Worms were fed with *lmn-1* (RNAi) or EV for 72h. (D) Relative length of muscle cells in N2 and *eat-2* worms, both expressing *myo-2*;RFP, that were fed with *lmn-1* (RNAi) or EV for 72h. *n*(N2) = 24, *n*(eat*-2*) = 26 worms and totals of 183 and 251 cells respectively were used for the analysis. *P*=7.37×10^−15^. (*E*) Average pumping rate of *eat-2* (*ad1116*) animals fed with *lmn-1* (RNAi) or EV for 72h. *n*=51. Error bars in all graphs represent mean ± SEM.

### Lamin regulates the reduced fat levels of DR animals

Animals subjected to DR accumulate less fat. This is due to a genetically controlled process, where nutrients are allocated elsewhere (Bar et al., 2016; Palgunow et al., 2012). To test whether *lmn-1* regulates fat accumulation in *C. elegans*, we measured fat levels using Oil Red O staining. Downregulation of *lmn-1* significantly increased fat levels in *eat-2* but not in control animals (**Fig. 2*A*** and ***B***, *p*<*0*.*0001*). These results indicate a role for *lmn-1* in regulating fat levels.

**Figure 2.**
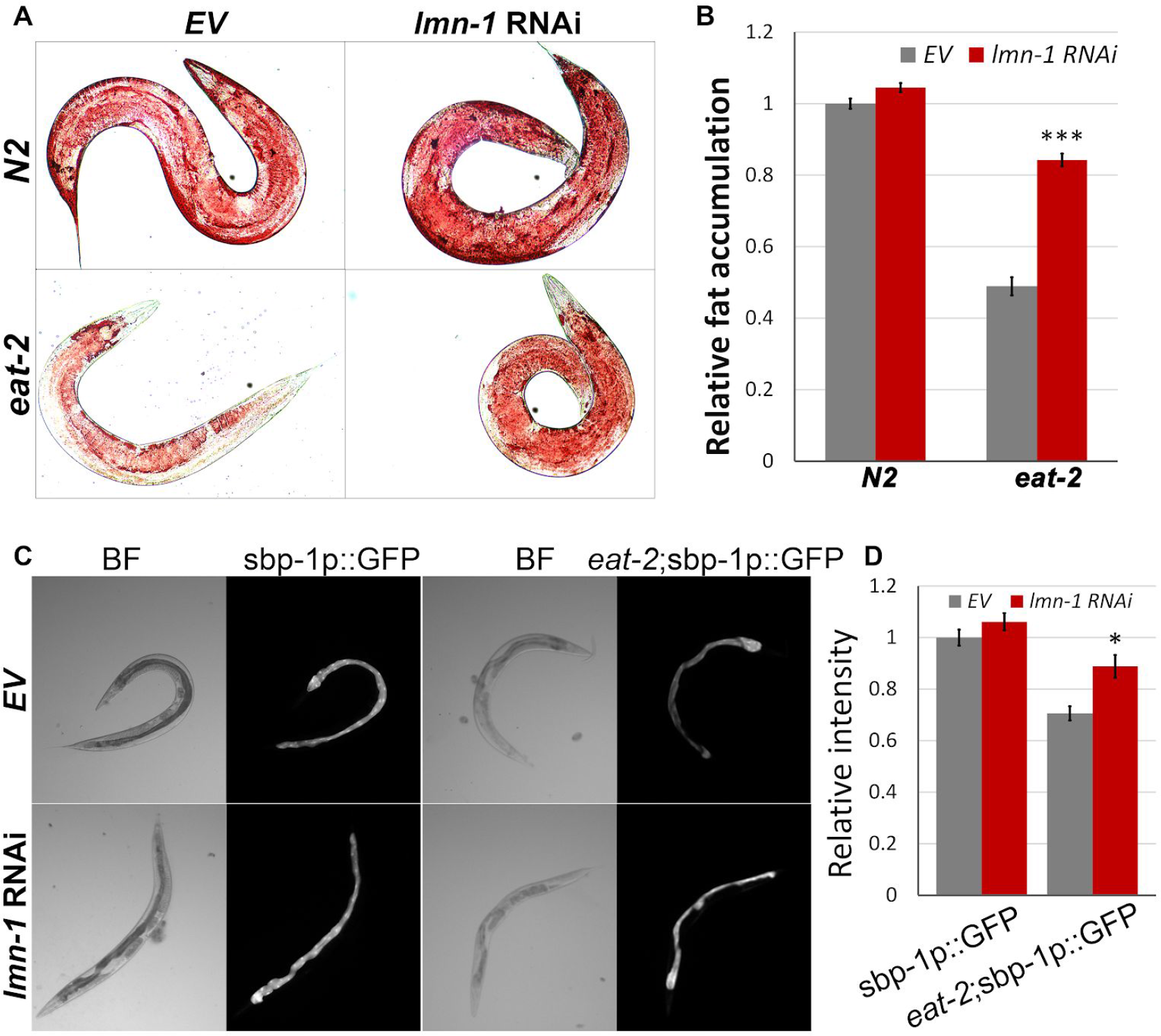
*lmn-1* regulates fat accumulation and *sbp-1* transcription in DR animals. (*A*) Representative stereomicroscope images of Oil Red O staining of control (N2) and *eat-2* (*ad1116*) worms fed with *lmn-1* (RNAi) or EV for 72h. (*B*) Average levels of relative intensity of Oil Red O staining in *eat-2* (ad1116) and in control (N2) animals fed with *lmn-1* (RNAi) or EV for 72h. *n*(N2)=99, *n*(*eat-2*)=75. (*C*) Representative fluorescence microscopy images of control (N2) and *eat-2* (*ad1116*) animals expressing GFP driven by the *sbp-1* (sbp-1p::GFP) promoter and fed with either EV or *lmn-1* (RNAi). (*D*) Quantification of (*C*). *n*(N2)=52, n(*eat-2*)=60. Error bars in all graphs represent mean ± SEM.

### Lamin regulates the transcription of sterol binding protein 1 (*sbp-1*)

Sterol regulatory element-binding protein (SREBP) is a transcription factor required for fatty acid biosynthesis. Sterol Binding Protein 1 (SBP-1) is the *C. elegans* homolog and a positive regulator of lipid storage (Sato, 2010). To test whether lamin regulates *sbp-1* transcription, we downregulated lamin in control and *eat-2* animals expressing GFP fused to the *sbp-1* promoter (sbp-1p::GFP). While *sbp-1* transcription levels are relatively low in *eat-2* animals (*p*<0.0001, **Fig. 2*C*** and ***D***), downregulation of *lmn-1* re-elevated *sbp-1* levels, most notably in the gut (**Fig. 2C** and **D**, *p*<*0*.*01*). We note that lamin is known to regulate SREBP subcellular localization (Duband-Goulet et al., 2011) and inhibition of mTOR causes SREBP to accumulate at the nuclear envelope, where it is inactive (Peterson et al., 2011). Thus, it is likely that the transcriptional regulation of *sbp-1* only partially accounts for excess fat accumulation in these DR animals.

### *lmn-1* regulates animal size upstream to the TORC1 complex

*atx-2* and *gdi-1* are the functional homologs of the Tuberous Sclerosis Complex (TSC) components in *C. elegans*, which decrease cell size via the inhibition of mTOR pathway (Bar et al., 2016). As both *lmn-1* and *atx-2* RNAi reverse the DR-induced size reduction (**Fig. 1*A*** and **3*A***), we mapped the relationship between these genes. When *eat-2* animals were subjected to RNAi against both genes, no additive increase in size was observed (**Fig. 3*A***). This suggests that *atx-2* and *lmn-1* share the same pathway, or at least the same downstream targets, with respect to body size regulation.

**Figure 3.**
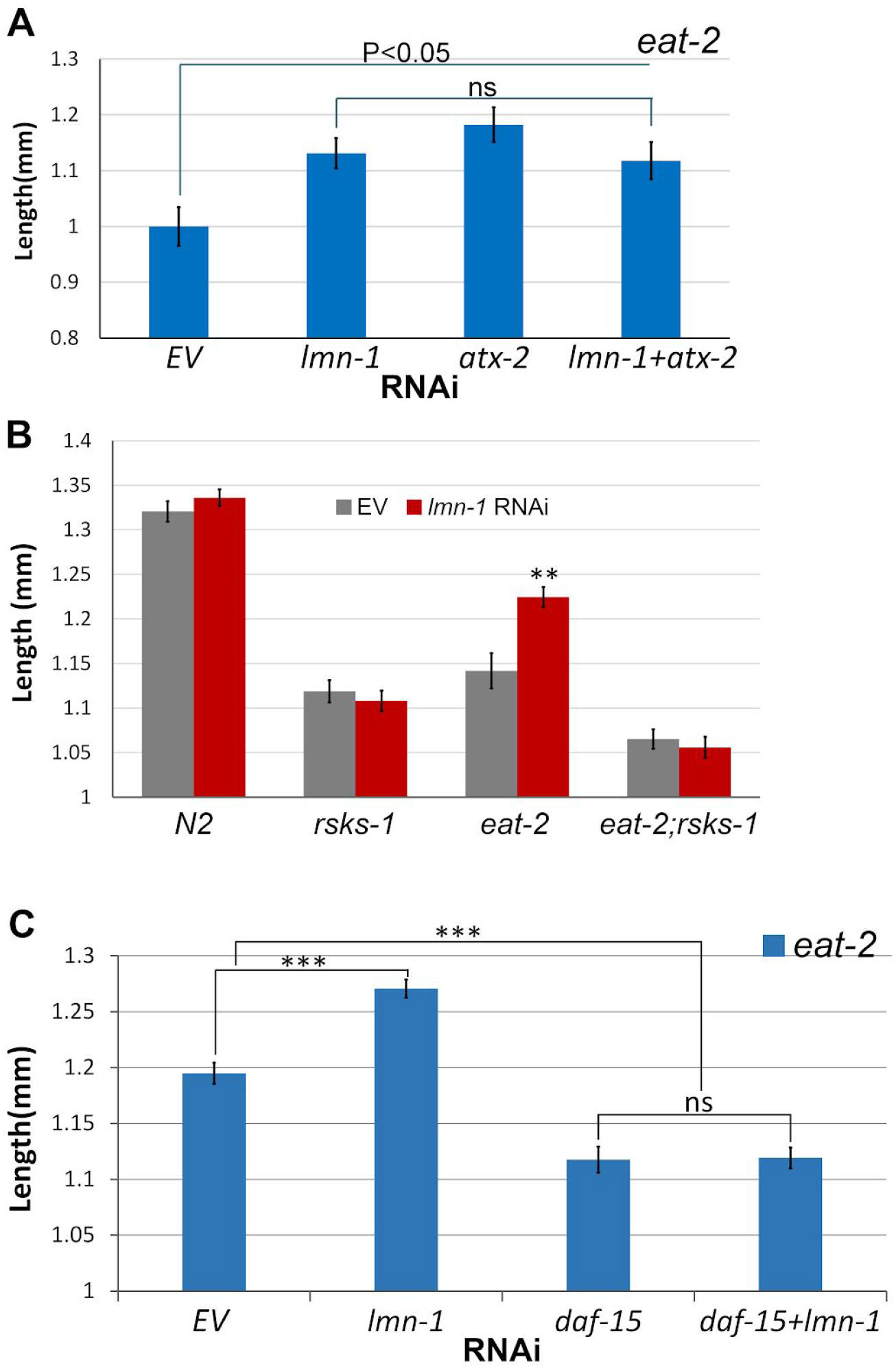
*lmn-1* regulates animal size upstream to the mTOR pathway. (*A*) Average length of *eat-2* young adult worms fed with RNAi of either *lmn-1, atx-2*, both *lmn-1* and *atx-2* or EV for 72h. (*B*) Average length of N2, *eat-2, rsk-1* or *eat-2;rsk-1* young adult worms fed with RNAi for *lmn-1* or EV for 72h. *n*(N2)=66, n(*rsks-1*)=67, *n*(*eat-2*)=59, n(*eat-2;rsks-1*)=54 (*C*) Average length of *eat-2* young adult worms fed with RNAi for either *lmn-1, daf-15*, both *lmn-1* and *daf-15* or EV. *n*(*eat-2*)=186. Error bars in all graphs represent mean ± SEM.

A*TX-2* regulates the mTOR pathway, which is dysregulated in some laminopathies. We further investigated the role of lamin in the mTOR pathway. Ribosomal protein s6 kinase beta-1 (S6K), is a downstream target of mTORC1 (Sfakianos et al., 2018). Its main activity is the phosphorylation of the s6 ribosomal protein, leading to an increase in protein synthesis and cell proliferation. Deletion of *rsks-1*, the homologue of S6K in *C. elegans*, resulted in a major decrease in the size of both *eat-2* and control worms (**Fig. 3*B***, *p*<0.0001 and *p*<0.01 respectively). This size decrease could not be rescued by downregulation of *lmn-1*, indicating that lamin regulates size upstream to S6K (**Fig. 3*B***). To confirm that the size increase due to lamin knockdown depends directly on mTOR itself, we downregulated *lmn-1* together with *raptor*, a key member of the mTOR complex-1 (mTORC1) (Carriere et al., 2011). Down-regulation of *daf-15*, the *C. elegans* homolog of *raptor*, decreased the size of *eat-2* animals (**Fig. 3*C***, *p*<0.0001) and eliminated the *lmn-1* (RNAi) size increase (**Fig. 3*C***). Based on these results we concluded that lamin acts upstream to the mTOR complex-1 to regulate size.

Under nutrient deprivation, the GDI-1:ATX-2 axis acts upstream to Ras homolog enriched in brain (RHEB), to change it from RHEB-Guanosine-5’-triphosphate (GTP) to RHEB-guanosine diphosphate (GDP). While RHEB-GTP promotes activation of mTORC1 complex, RHEB-GDP blocks its activity (Bar et al., 2016; Laplante and Sabatini, 2012). This leads to an opposite effect on fed (N2) animals, decreased in size due to elimination of RHEB-GTP and reduced activation of mTOR, and increased size in DR animals, due to elimination of RHEB-GDP and reduced inhibition of mTOR (Bar et al., 2016). Indeed, down regulation of *rheb-1* in control animals decreased their size (**Fig. 4*A***, *p*<*0*.*0001*), however down regulation of *lmn-1* reversed this effect (**Fig. 4*A***, *p*<*0*.*0001*). Under DR conditions, downregulation of *rheb-1* increases their size (**Fig. 4*B***, *p*<*0*.*0001*), while downregulation of both *lmn-1* and *rheb-1* had no additive effect (**Fig. 4*B***). We concluded that lamin might function downstream to RHEB in regulating the mTOR pathway.

**Figure 4.**
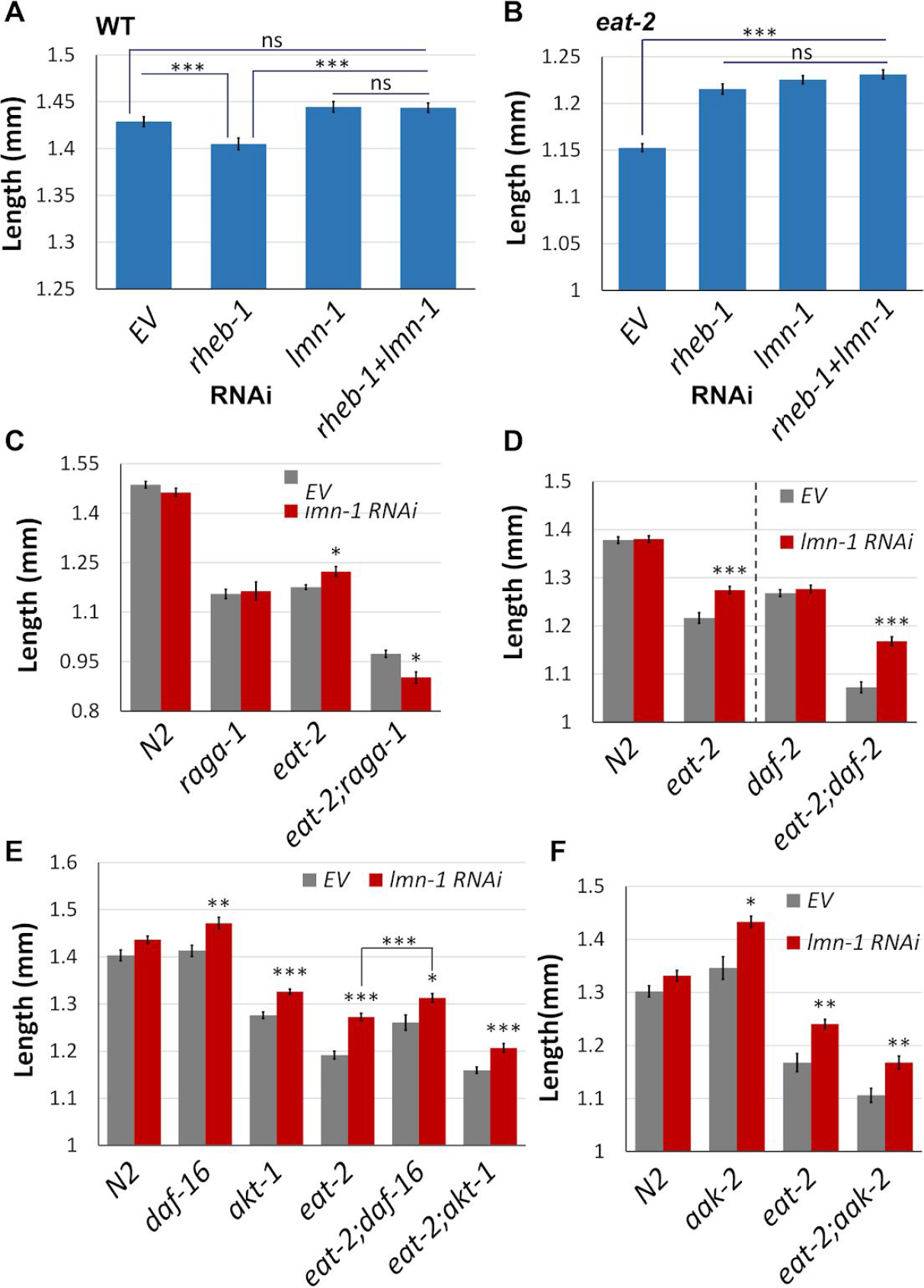
Mapping *lmn-1* to the mTOR pathway. (*A*) Average length of N2 young adult worms fed with RNAi for either *lmn-1, rheb-1*, both *lmn-1* and *rheb-1* or EV. *n*(N2)=494. (*B*) Average length of *eat-2* young adult worms fed with RNAi for either *lmn-1, rheb-1*, both *lmn-1* and *rheb-1* or EV. *n*(*eat-2*)=680 (*C*) Average length of N2, *eat-2, raga-1* or *eat-2;raga-1* young adult worms fed with *lmn-1* (RNAi) or EV. *n*(N2)=70, n(*raga-1*)=92, *n*(*eat-2*)=73, n(*eat-2;raga-1*)=45 (*D*) Average length of N2, *eat-2, daf-2* or *eat-2;daf-2* young adult worms fed with *lmn-1* (RNAi) or EV. *n*(N2)=96, n(*eat-2*)=92, *n*(*daf-2*)=72, n(*eat-2;daf-2*)=51 (*E*) Average length of N2, *eat-2, daf-16, akt-1, eat-2;daf-16* or *eat-2;* young adult worms fed with l*mn-1* (RNAi) or EV. *n*(N2)=80, n(*daf-16*)=95, n(*akt-1*)=115, *n*(*eat-2*)=72, n(*eat-2;daf-16*)=82, n(*eat-2;akt-1*)=97 (*F*) Average length of N2, *eat-2* or *aak-2* young adult worms fed with *lmn-1* (RNAi) or EV. *n*(N2)=50, n(*aak-2*)=43, *n*(*eat-2*)=42, n(*eat-2;aak-2*)=57. Error bars in all graphs represent mean ± SEM.

A key regulator of the mTOR pathway is the Ras-related GTP-binding protein (RAG) complex. Dependent on amino acids availability, the subunits RAG(A/B)-GTP and RAG(C/D)-GDP form a heterodimer that recruits and anchor mTOR to the lysosome (Sancak et al., 2008; Wu et al., 2016), where it can be activated by its upstream regulators. Deletion of *raga-1*, the homologue of RAG(A/B), decreases the size of control animals, and DR conditions further decrease their size (**Fig. 4*C***, *p*<*0*.*001*). Down regulation of *lmn-1* in DR worms lacking *raga-1* resulted in additional size reduction (**Fig. 4*C***), suggesting that *raga-1* is essential for *lmn-1* activity in the context of size regulation. The additional size decrease may suggest that *lmn-1* has mTOR-independent roles that may affect size.

### *lmn-1* size regulation is not dependent on the insulin/IGF-1 signaling pathway

Growth factors stimulate cellular growth and proliferation partially via the mTOR pathway. Activated by the growth factors, insulin growth factor receptors (IGFRs) set a phosphorylation cascade that eventually promotes mTOR activity (Inoki et al., 2003; Laplante and Sabatini, 2012). *Daf-2* is the sole member of the IGFR family in *C. elegans*. Down regulation of *lmn-1* in *eat-2* worms, also mutated in *daf-2*, resulted in an increase to their size (**Fig. 4*D***, *p*<*0*.*0001*), suggesting that *lmn-1* is not dependent on *daf-2* in size regulation of *eat-2* animals. AKT-1 is triggered by IGF signaling (Laplante and Sabatini, 2012). Once activated, AKT-1 activates the mTOR pathway by inhibiting phosphorylations of the TSC (Inoki et al., 2002; Manning et al., 2002). As expected, deletion of *akt-1* resulted in size decrease both in control and *eat-2* animals (**Fig. 4*E***, *p*<*0*.*0001* and *p*<*0*.*01*, respectively). However, downregulation of *lmn-1* increased animal size both in control and *eat-2* animals (**Fig. 4*E***. *p*<*0*.*0001*), indicating that *akt-1* is not required for *lmn-1* size regulation.

Adenosine monophosphate-activated protein kinase (AMPK), is a master regulator of cellular energy levels (Inoki et al., 2003). Downregulation of *lmn-1* in *eat-2* worms also lacking *aak-2*, the catalytic subunit of AMPK, increased their size (**Fig. 4*F***, *p*<*0*.*001*). *DAF-16*, a target of AMPK and DAF-2, is a transcription factor that translocates to the nucleus upon multipole stress signals, and inhibits mTORC1 (Robida-Stubbs et al., 2012). Deletion of *daf-16* resulted in a size increase of *eat-2* animals (**Fig. 4*E***, *p*<*0*.*001*). Downregulation of *lmn-1* in *eat-2* animals lacking *daf-16* resulted in an additional size increase (**Fig. 4*E***, *p*<*0*.*01*). This suggests that *daf-16* and *lmn-1* regulate the mTOR pathway at least partially independently.

### Dietary restriction delays age dependent nuclear envelope deformation

The nuclear lamina acquires structural deformation, including lobulations, membranes invagination and aggregations, in an age dependent manner (Haithcock et al., 2005). To test the effect of dietary restriction on the nuclear lamina, we imaged animals expressing LMN-1::GFP driven by the *lmn-1* promoter at day 2, 4 and 6 of adulthood, and classified the nuclear shape into 3 groups based on deformation severity (**Fig. 5*A*-*G***). At day 2 of adulthood, control animals began to show mild ageing phenotype, as more than 30% of the nuclei demonstrated mild abnormalities such as membrane folding and some lamin foci (class II) (**Fig. 5*A*** and ***G***). At day 4 of adulthood this phenotype was aggravated, as animals showed increased lobulations and lamin aggregations (45% class II and III; **Fig. 5*B*** and ***G***). This was further aggravated at day 6 as more control animals accumulated nuclear mis-shapes, such as stretched nuclei, nuclear lobulations and increased membrane folding. Overall, more than 50% of the nuclei shifted toward class II and III (**Fig. 5*C*** and ***G***). By contrast, DR animals showed a lag in nuclear envelope ageing phenotypes. At day 2, most of the nuclei (∼93%) did not exhibit any nuclear deformation as the nuclei were smooth and round (class I), with only a small fraction showing nuclear abnormalities (∼7%) (**Fig. 5*D*** and ***G***). Moreover, at day 4 and even 6 of adulthood, most of the nuclei still kept their un-deformed shape (86% and 82%, respectively at class I; **Fig. 5*E, F*** and ***G***). These results were accompanied by a decrease in total LMN-1 levels from day 2 to day 4 in control animals, and a smaller decrease from day 4 to day 6 in DR animals (**Fig. 5*H*** and ***J***). Based on these results we concluded that DR conditions delay age dependent nuclear deformation, as DR nuclei retain their smooth shape and structure even at advanced age. Some DR phenotypes are dependent on AMPK and its downstream targets (**Supp. Fig. 2** and (Uno and Nishida, 2016)). We tested whether the protective effects of DR on the nuclear morphology depends on this pathway as well. Downregulation of *aak-2* resulted in a faster accumulation of nuclear abnormalities (**Supp. Fig. 2**), suggesting this is a regulated process.

**Figure 5.**
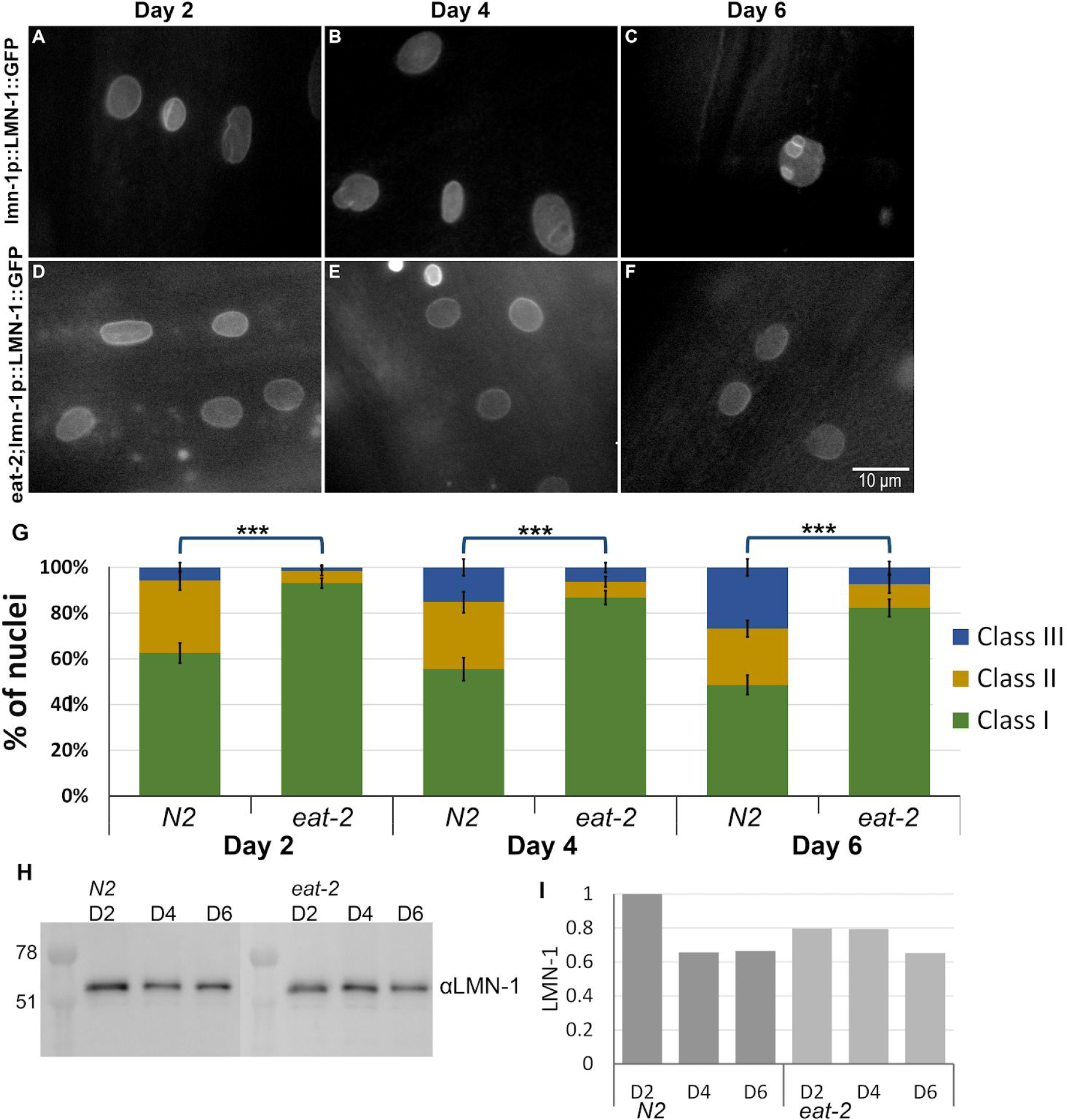
Dietary restrictions delay age dependent nuclear deformation. (*A*-*F*) Representative microscope images of control (N2) (*A*-*C*) and *eat-2* (*D*-*F*) worms expressing LMN-1::GFP at days 2 (*A,D*), 4 (*B,E*) and 6 (*C,F*) of adulthood. (*G*) Relative distribution of the 3 different classes grading (see materials and methods) the nuclear morphology changes in N2 and *eat-2* animals at days 2, 4 and 6 of adulthood. n(D2)=255 nuclei, n(D4)=228 nuclei, n(D6)=238 nuclei. Note: Error bars in all graphs represent mean ± SEM. *P* values were calculated using Fisher Exact Probability Test. (*H*) Western blot gel image of LMN-1 protein in N2 and *eat-2* at days 2, 4 or 6 of adulthood. (*I*) LMN-1 levels normalized according to total protein count and nematode number.

### Dysregulation of mTOR drives nuclear envelope deformation

Animals with life-extending mutations can, under specific conditions, show a delayed accumulation of nuclear envelope abnormalities ((Haithcock et al., 2005; Pérez-Jiménez et al., 2014) and **Fig. 6*A*-*D*** and ***I***). By contrast, laminopatic mutations can drive the premature accumulation of such defects (van Tienen et al., 2019).

**Figure 6.**
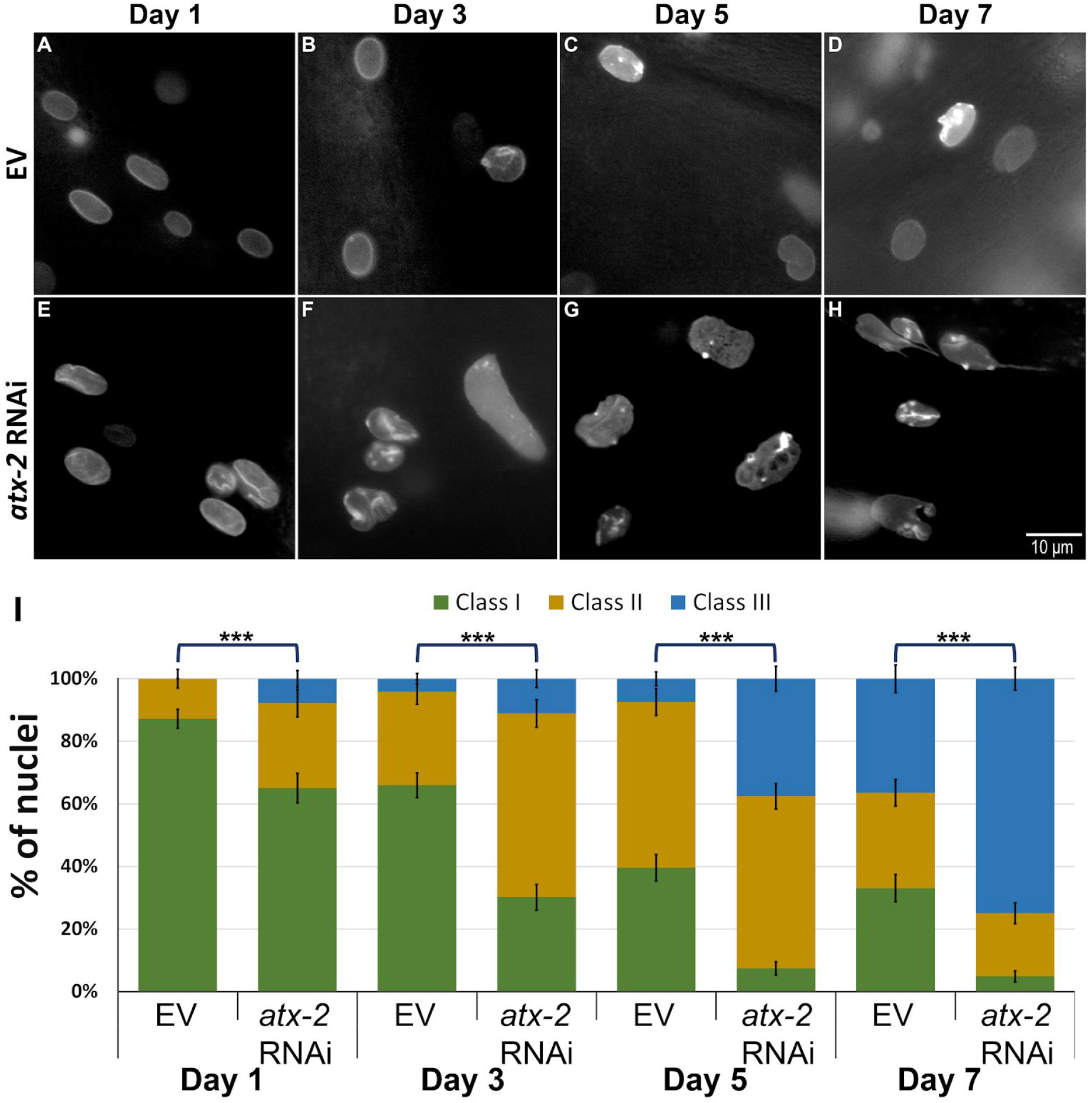
ATX-2 is required to maintain nuclear structure. (*A*-*H*) Representative microscope images of worms expressing LMN-1::GFP fed by either EV (*A*-*D*) or *atx-2* (RNAi) post-hatching (*E*-*H*), at day 1 (*A,E*), day 3 (*B,F*), day 5 (*C,G*) and day 7 (*D,H*) of adulthood. (*I*) Relative distribution of the 3 different classes grading the nuclear morphology changes in LMN-1::GFP expressing worms fed with *atx-2* (RNAi) or EV at days 1, 3, 5 and 7 of adulthood. n(D1)=228 nuclei, n(D3)=267 nuclei, n(D5)=283 nuclei, n(D7)=262 nuclei. Error bars in all graphs represent mean ± SEM. P value was calculated using Fisher Exact Probability Test.

When *atx-2* was knocked-down in young adult animals, we saw a small but statistically significant increase in nuclei showing abnormalities. To aggravate this phenotype and to test the role of *atx-2* in nuclear shape regulation, we subjected animals expressing LMN-1::GFP, driven by the *lmn-1* promoter, to RNAi downregulation of *atx-2* immediately post-hatching. Changes in nuclear morphology were documented every other day, starting at day 1 of adulthood and until day 7. Nuclear morphology was classified into 3 class groups as described in the previous section. Already at day 1 of adulthood, nuclei of *atx-2* (RNAi) adult animals showed increased lobulations and lamin aggregations at the nuclear envelope (**Fig. 6*E*** and ***I***). These age-related phenotypes were much more robust at day 3, as more nuclei shifted to class II and class III (**Fig. 6*F*** and ***I***). By day 5, only 7% of the nuclei were smooth (class I) (**Fig. 6*G*** and ***I***). By day 7 of adulthood, 75% of the nuclei were fragmented, stretched, showed invagination of the membrane and severe lobulation (class III) while only a small fraction were in class I (**Fig. 6*H*** and ***I***). In line with these findings, accelerated breakdown of the nuclear envelope following *atx-2* downregulation was observed in transgenic worms expressing the nuclear envelope protein *emr-1*, fused to GFP (**Supp Fig. 3**). We concluded that the mTOR pathway regulates lamin distribution, and that downregulation of *atx-2* results in nuclear envelope phenotypes, similar to those seen in older animals (Haithcock et al., 2005).

### *lmn-1* regulates RAGC-1 localization to the nucleus

To anchor mTORC1 to the lysosome, RAGC has to enter the nucleus as RAGC-GTP. In the nucleus, RAGC acquires its active RAGC-GDP form and exits back to the cytoplasm (Wu et al., 2016) where it anchors mTORC1 to the lysosome. To follow the expression and localization of RAGC-1, the *C. elegans* homologue of RAG (C/D), we used CRISPR/Cas9 to fuse a FLAG-tag to the endogenous RAGC-1 (**Fig. 7*A***). While in control animals RAGC-1 was expressed both in the cytoplasm and the nucleus (**Fig. 7*B***), in *eat-2* animals, RAGC-1 was mostly excluded from the nucleus (**Fig. 7*C*** EV panel). We compared the ratio of cytoplasmic RAGC-1 to nuclear RAGC-1 (**Fig. 7*D***). In control animals, as well as control animals subjected to *lmn-1* RNAi, this ratio was close to one, indicating equal presence of RAGC-1 in the nuclear and in the cytoplasmic fractions. By contrast, in DR worms, this ratio was at least 3 times higher than in control animals, possibly indicating a limited shuttling of RAGC-1. If lamin regulates the mTOR pathway through RAGC-1 localization, we would expect downregulation of *lmn-1* to restore it to the nucleus. Indeed, following *lmn-1* (RNAi), RAGC-1 entered the nucleus, potentially allowing it to acquire its active form (**Fig. 7*C*** and ***D***).

**Figure 7.**
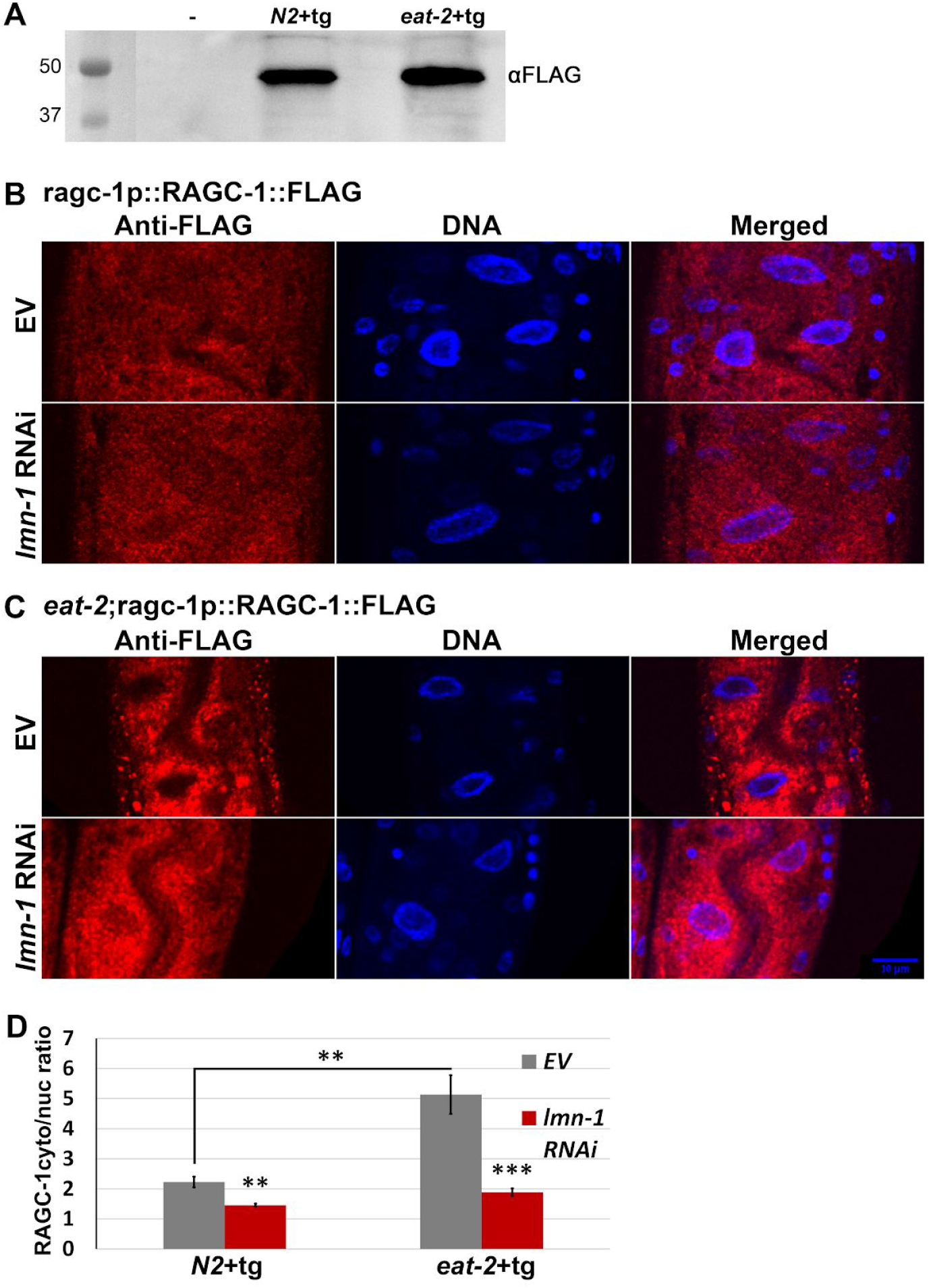
*lmn-1* regulates the entrance of RAGC-1 to the nucleus. (*A*) Western blot of transgenic ragc-1p::RAGC-1::FLAG in N2 and *eat-2* worms. (*B*-*C*) Representative confocal images of N2 (*B*) and *eat-2* (*C*) animals expressing RAGC-1 fused to FLAG-tag (RAGC-1::FLAG) that were fed with *lmn-1* (RNAi) or EV for 48h post-hatching (*D*) Relative distribution (cytoplasmic to nuclear) of RAGC-1 in N2 and *eat-2* worms expressing RAGC-1 fused to a flag tag. *n*(N2)=19, *n*(*eat-2*)=29 worms and totals of 50 and 65 cells, respectively, were used for the analysis. Error bars in all graphs represent mean ± SEM.

## Discussion

### Lamin regulates DR

Mutations in *LMNA* genes are associated with ageing disorders as well as metabolic diseases. One example is Hutchinson-Gilford progeria syndrome (HGPS), where lamin mutations result in an accelerated ageing disorder and pathological phenotype (Goldman et al., 2004; Gonzalo et al., 2017). A recent study in HGPS mouse models shows the potential of DR in improving life expectancy (Bárcena et al., 2018). Here we showed that in DR *C. elegans*, multiple DR related phenotypes are regulated by lamin. Downregulation of *lmn-1* in DR animals partially rescues their size and fat levels, while having minimal effect on control animals. Two other aspects of DR, fecundity and longevity, could not be properly analysed in a similar way, as lamin is essential for development and its downregulation also significantly shortens lifespan of both control (N2) and DR animals (Bar et al., 2016).

### Lamin acts through the mTOR pathway

Multiple studies have shown a cross talk between lamin genes and the mTOR pathway (Chiarini et al., 2019; Lattanzi et al., 2014). The mTOR pathway is affected by some *LMNA* mutations, and mTOR inhibitors are successfully used to treat multiple laminopatic mutations in cell lines, in animal models and recently in clinical trials (Pellegrini et al., 2015; Ramos et al., 2012). We explored the regulatory role of *lmn-1* in DR using an epistasis assay. Three genes were found to be required for *lmn-1* (RNAi) size rescue: S6K, a target of the mTOR pathway; *RAPTOR*, a core protein of the mTORC1 complex; and RAG(A/B), an mTOR regulator. We concluded that mTORC1 is essential for *lmn-1* size dependent regulation.

### ATX-2 regulates nuclear morphology

As cells age, their nuclei accumulate abnormalities (Haithcock et al., 2005). DR animals accumulated abnormalities slower than control animals (**Fig. 5*A*-*G***). Activation of the mTOR pathway, by downregulation of the negative regulator *atx-2*, resulted in a significant increase in nuclear abnormalities (**Fig. 6**). In mammalian cells, inhibition of mTORC1 affects the nuclear shape through regulation of lipin 1 nuclear localization (Peterson et al., 2011). One possible mechanism of nuclear shape regulation is via post-translational modifications of lamins. Phosphorylation of lamins determines their structural properties and affects signalling (Torvaldson et al., 2015). Insulin, a known mTOR activator, drives lamin phosphorylation during interphase (Friedman and Ken, 1988). It will be interesting to test if post-translational modifications to lamins and other nuclear envelope proteins may regulate the mTOR pathway under DR.

### RAGC-1 exclusion from the nucleus in DR requires an intact nuclear lamina

Lamin Directly interacts with the Linker of Nucleoskeleton and Cytoskeleton (LINC) complex and the Nuclear Pore Complex (NPC) (Bone et al., 2014; Guo and Zheng, 2015). The NPC regulates nuclear transport. Several laminopathic mutations disrupt nucleocytoplasmic translocation (Busch et al., 2009; Han et al., 2019; Kelley et al., 2011). Some laminopathies are characterized by failure to exclude proteins from the nucleus. Yes-Associated Protein (YAP) is a sensor and mediator of mechanical cues. Mutations resulting in *LMNA* related congenital muscular dystrophy cause the NPC to fail to exclude YAP from the nucleus, contributing to the disease pathology (Owens et al., 2020). A morphological hallmark of the different laminopathies is a deformed nuclear envelope (van Tienen et al., 2019). One plausible mechanism for the impaired nuclear transport in laminopathies is that the altered nuclear shape causes changes of the stretch forces applied on the NPC, a known regulator of the flux (Donnaloja et al., 2019). Nuclear deformations are also found in aged nuclei (Haithcock et al., 2005; Pérez-Jiménez et al., 2014), along with compromised nucleocytoplasmic transportation (D’Angelo et al., 2009). Recently, it was shown that nuclei of aged *C. elegans* are more susceptible to damaging effects of outside stretch forces (Zuela-Sopilniak et al., 2020). This is likely the result of altered nuclear stiffness and lower elasticity, suggesting that aged nuclei are under constant self-induced stretch. A similar phenomena is observed when the LINC complex is impaired (Zuela-Sopilniak et al., 2020). When nuclei are exposed to outside forces that deform the nuclear envelope morphology, NPC are redistributed and stretched, increasing transcription factor nuclear import (Elosegui-Artola et al., 2017; Hoffman et al., 2020). In mammalian cells, nuclear eccentricity is regulated by nutrients and mTORC1. In *C. elegans*, DR delays age related nuclear morphology phenotypes and excludes RAGC-1 from the nucleus, thus inhibiting mTOR. On the other hand, compromising the nuclear envelope by downregulation of *lmn-1* restores RAGC-1 into the nucleus, activates mTOR and increases animal size and fat content. This model also explains the increase seen in *sbp-1* transcription, known to be regulated by mTOR (Bar et al., 2016; Li et al., 2011; Peterson et al., 2011).

Downregulation of lamin likely affects multiple genes and proteins, as the nuclear envelope preferentially interacts with both DNA lamina associated domains (LADs) and multiple proteins, including transcription factors and chromatin remodelers. However, we found that in DR animals, the size effect is completely dependent on the RSKS-1, DAF-15 and RAGC-1. One interpretation of these data is that lamin, via the NPC and nucleocytoplasmic transport, regulates the mTOR pathway. Under DR, and potentially other types of stress, the nuclear lamina is required to make the NPC less permissive. Stress-dependent post-translational modifications may drive these changes. The less permissive NPC prevents passive nuclear entranry of RAGC-1, and potentially other proteins. A similar mechanism was demonstrated for biguanides, that regulate the mTOR pathway by preventing RAGC from passing through the NPC (Wu et al., 2016). These findings may have implications on our understanding of how the nuclear lamina regulates gene expression in other situations, including mechanotransduction. Future studies will test whether laminopathic mutations affect the permissibility of the NPC, and whether it is linked to their pathologies.

## Materials and methods

### *C. elegans* strains

Strain maintenance and manipulations were performed under standard conditions as previously described (Brenner 1974). All experiments were performed at 23°C unless described otherwise. The following strains were used: N2 (control); DA116, *eat-2*(*ad1116*) II; CF1038, *daf-16*(*mu86*) I; YG2227, (*eat-2*(*ad1116*) II; *daf-16*(*mu86*) I); RB754, *aak-2*(*ok524*) X; YG2223, (*eat-2*(*ad1116*) II; *aak-2*(*ok524*) X); RB1206, *rsks-1*(*ok1255*) III; YG2229,(*eat-2*(*ad1116*) II; *rsks-1*(*1255*) III); CF1041, *daf-2*(*e1370*); YG2606,(*eat-2*(*ad1116*) II; *daf-2*(*e1370*)); RB759, *akt-1*(*ok525*)*V;* YG2605, (*eat-2*(*ad1116*) II; *akt-1*(*ok525*)*V*); *VC222*, [raga-1(*ok386*)](https://cgc.umn.edu/gene/246558) *II;* (*eat-2*(*ad1116*) II; *VC222*, [raga-1(*ok386*)](https://cgc.umn.edu/variation/1896130) *II*); [*sbp-1p::GFP*]; *YG2604*, (*eat-2*(*ad1116*) II; [sbp-1p::GFP]); YG2607,[ragc-1p::RAGC-1::FLAG]; YG2608,((*eat-2*(*ad1116*); YG2607[ragc-1p::RAGC-1::FLAG]); PD4810 (*lmn-1*:GFP, *ccIs4810* [pJKL380.4 P*lmn-1*::*lmn-1*::gfp::*lmn-1* 3’UTR +myo-2p::MYO-2::RFP]; LW699,[*lmn-1p*::*lmn-1*::gfp::unc-54 3’UTR+ unc-119(+)]. All strains were obtained from the *C. elegans* Genome Center (CGC), generated using micro injection or obtained using genetic crossing.

### Generating *ragc-1*p::RAGC-1::FLAG strain using CRISPR-Cas9

ragc-1p::RAGC-1::FLAG, was generated by microinjection using Crispr-Cas9 as described previously (Friedland et al. 2013), and verified by sequencing and western blotting. A 3X FLAG tag was inserted to the *ragc-1* sequence after the ATG start codon as follows:

### crRNA-

AATCATCAAAATCCTCGTCA

### ssODN-

**Start codon** modified base to cancel NGG **<u>crRNA</u> <u>site</u> *<u>Restriction</u> <u>site</u>*** (3X FLAG is in UPPERCASE) taaaccttttcaatttcaga**atg**GACTACAAGGACCACGACGGAGACTACAAGGACCACGA TATCGATTACAAGGACGACGACGACAAGgagtcgga**t**cc**tg*ac*gaggattttgatgatt**accgct

### Statistics

Unless noted otherwise, comparisons were performed using an unpaired two-tailed Student *t* test. A minimum of 3 biological repeats per experiment were performed.

### RNAi experiments

RNAi feeding experiments were performed as described previously (Ahringer, 2006). In brief, nematode growth medium (NGM) plates containing 25 μg/mL ampicillin and 1mM isopropyl β-d-thiogalactoside (IPTG) were seeded with the appropriate bacteria taken from either the Ahringer library (Kamath and Ahringer, 2003) or the Vidal library (Rual et al., 2004). Controls were placed on feeding plates with an empty L4440 vector (EV). Worms were placed on plates at the appropriate developmental stage and analyzed for various phenotypes. When antibodies or GFP fusions were available, the down-regulation efficiency was verified by analysis of the proteins level. In other cases, the level of mRNAs of the perturbed gene was analyzed by quantitative Polymerase Chain Reaction, or the animal phenotypes were analyzed by matching the published data.

### Size experiments

Animals were synchronized by bleaching (Porta-de-la-Riva et al., 2012) or by allowing them to lay eggs on plates for 6h. In RNAi experiments, except for *daf-15* downregulation and *daf-2* mutants, animals were allowed to develop on NGM plates for 48h until the *eat-2* worms reached larval stage 3. Animals were transferred to RNAi plates, and then to fresh RNAi plates every 48h, to exclude progeny. Unless noted otherwise, size measurements were performed after 72h of growth on RNAi plates. In the *daf-15* downregulation assays, after bleaching, nematodes were placed on *daf-15* RNAi plates for 48h then moved to fresh *daf-15* or *daf-15 + lmn-1* plates. Animals carrying the *daf-2* (*e1370*) mutation were allowed to develop on NGM plates for 96h at 16°C, until *daf-2;eat-2* nematodes reached larval stage 3, then transferred to fresh RNAi plates for 72h at 23°C. Imaging was performed using Olympus MVX10 dissecting microscope with a Dino-Eye camera (AnMo Electronics) and animals’ length from head to tail was measured using ImageJ. Ten to fifty worms were measured at each time point. These experiments were repeated at least three times. To measure muscle cell length, transgenic animals containing MYO-2 protein fused to RFP were mounted on 2% agarose pads with 2mM levamisole. Images were acquired with ORCA-R2 camera (Hamamatsu Photonics) mounted on an Axioplan 2 microscope with X60 oil lens. Length was measured from vertex to vertex along the muscle cell using ImageJ. At least 20 worms were measured at each time point. *P* values were calculated using a two-tailed *t* test.

### Oil Red O Staining

Oil Red O staining and quantification were performed as described previously (O’Rourke et al., 2009). Images were acquired using Nikon Eclipse E200 microscope with an X4 or X10 lens fitted with a Moticam 2300 color camera (Motic). Exposure levels were adjusted for clear staining without saturation, and maintained for all samples. Quantification was performed with the ImageJ package. *P* values were calculated using a two-sample *t* test for unequal variances. Experiments were repeated three times.

### *sbp-1*p::GFP analysis

Animals expressing GFP driven by the *sbp-1* promoter of the *sbp-1* gene, were synchronized to the fourth larval stage. Animals were placed on feeding plates with *lmn-1* (RNAi) for 48h, then transferred to new RNAi feeding plates for additional 24h. Controls were placed on feeding plates with an empty L4440 vector (EV). Each experiment included ∼30 worms. Images were acquired using an Olympus MVX10 microscope with QImaging Photometric 5MP camera. Quantification of GFP fluorescence intensity was performed using the ImageJ package. *P* values were calculated using a two-sample *t* test for unequal variances.

### Immunofluorescence

Animals were synchronized by bleaching, eggs were then laid on *lmn-1* RNAi or EV (control) feeding plates. Worms were then allowed to develop for 48h until *eat-2* animals reached larval stage 4. Worms were washed from the plates using M9 and transferred to 1.5ml tubes. Worms were washed 3x with M9 and 3x with 1X phosphate buffered saline (PBS) containing 0.1% Tween 20 (PBST) to remove excess bacteria. Worms were fixed with freshly made 2% formaldehyde in PBST followed by snap freeze with liquid nitrogen and 10 min incubation at room temperature (RT). Samples were washed 3X with PBST and permeabilized using mild sonication (Sonics Vibra-Cell sonicator) with a model CV33 microtip at ∼25% amplification 2×8 sec sonication periods. Samples were incubated for 25 min in X2 Modified Ruvkun’s Witches Brew (MRWB) (“Gonad-intestine Staining Protocol,” n.d.) followed by 3X washes with PBST, and incubated for 12.5 min with 10mM DL-Dithiothreitol (DTT). Samples were washed 4X with PBST then incubated for 1h in PBST followed by 10 min treatment with 0.3% H_2_O_2_. Samples were then washed 3X with PBST and blocked for 1h with a blocking buffer containing 1% bovine serum albumin in PBST. After blocking, samples were incubated overnight at RT, shaking, with a primary mouse anti-FLAG antibody (clone M2; monoclonal; Sigma, CAT # F1804) diluted in blocking buffer (1:500). After 4x washes for ∼2h with PBST, samples were incubated with the secondary anti-mouse-Cy3 antibody diluted in a blocking buffer (1:100) for ∼2h at RT. After 4x washes in PBST for 2h, DNA was stained for 10 min at RT using 4′, 6-Diamidino-2-phenylindole dihydrochloride (DAPI; Sigma, Cat# 28718903) diluted in PBST (1:1000), followed by a 3x 10 min washes in PBST. Samples were mounted on glass slides in a drop of mounting media (Vectashield, CAT# H-1400) and sealed with nail polish. Staining quantification was performed using the ImageJ package. *P* values were calculated using a two-sample *T-*test for unequal variances.

### Abnormal nuclear morphology classification

Transgenic control (N2) and *eat-2* animals expressing GFP fused to *lmn-1* driven by the promoter of the *lmn-1* gene and control animals expressing *emr-1* fused to GFP under *lmn-1* promoter, were synchronized by bleaching. Embryos were placed on feeding plates containing either EV, *atx-2* or *aak-2* RNAi. Worms were moved to fresh RNAi plates every 48h. Images were acquired at days 2, 4 and 6 (**Figs. 6** and **7**) or at days 1, 3, 5 and 7 (**Fig. 7**) of adulthood. Nuclei were counted and grouped into three different classes (class I–III) based on their morphology: class I - nuclei without any nuclear deformation (smooth nuclei); class II-nuclei with mild nuclear abnormalities such as membrane folding and some lamin foci; class III-nuclei with severe nuclear deformation including increased nuclear folding, lobulations and abnormal nuclei shape. Nuclei were counted from the middle part of the worm (excluding head, tail and gonads) and of either hypodermal or muscular origin; neuronal nuclei were excluded. For each time point 8-11 animals were used. *P* value was calculated using Fisher Exact Probability Test.

### Microscopy

Image 1A was obtained using Dino-Eye camera mounted on an Olympus MVX10 dissecting microscope. Image **1*C*** was obtained using ORCA-R2 camera (Hamamatsu Photonics) mounted on an Axioplan 2 microscope with X20 magnification. Images in **Fig. 2*A*** were obtained using Nikon Eclipse E200 microscope with an X4 or X10 magnifications fitted with a Moticam 2300 color camera (Motic). Images in 2*C* (quantification of GFP expression) were recorded using an Olympus MVX10 microscope. Images in **Figs. 5** and **6** were obtained using ORCA-R2 camera (Hamamatsu Photonics) mounted on an Axioplan 2 microscope with X100 magnification. Images in **Fig. 7*B*** and ***C***, were obtained using Leica SP5 confocal microscope. Panels ***A*-*F*** of **Fig. 5** were obtained at the same exposure, but image levels were adjusted individually to improve feature visibility, as was the case of panels ***A*-*H*** of **Fig. 6**. The internal panels of **Fig. 7*B*** are displayed under identical conditions, as are the internal panels of **7*C***. However, the image levels of **7*B*** and **7*C*** are not identical.

## Acknowledgments

We thank Prof. Michal Goldberg for a critical reading of the manuscript, Gabriel Bonduryansky, Naama Zung and the lab of Yonatan Tzur for technical assistance and suggestions, specifically Hanna Achache and Yisrael Rappaport. We also thank the Israeli Science Foundation (grant 632/20).

## Supplementary figures

**Supplementary figure 1.**
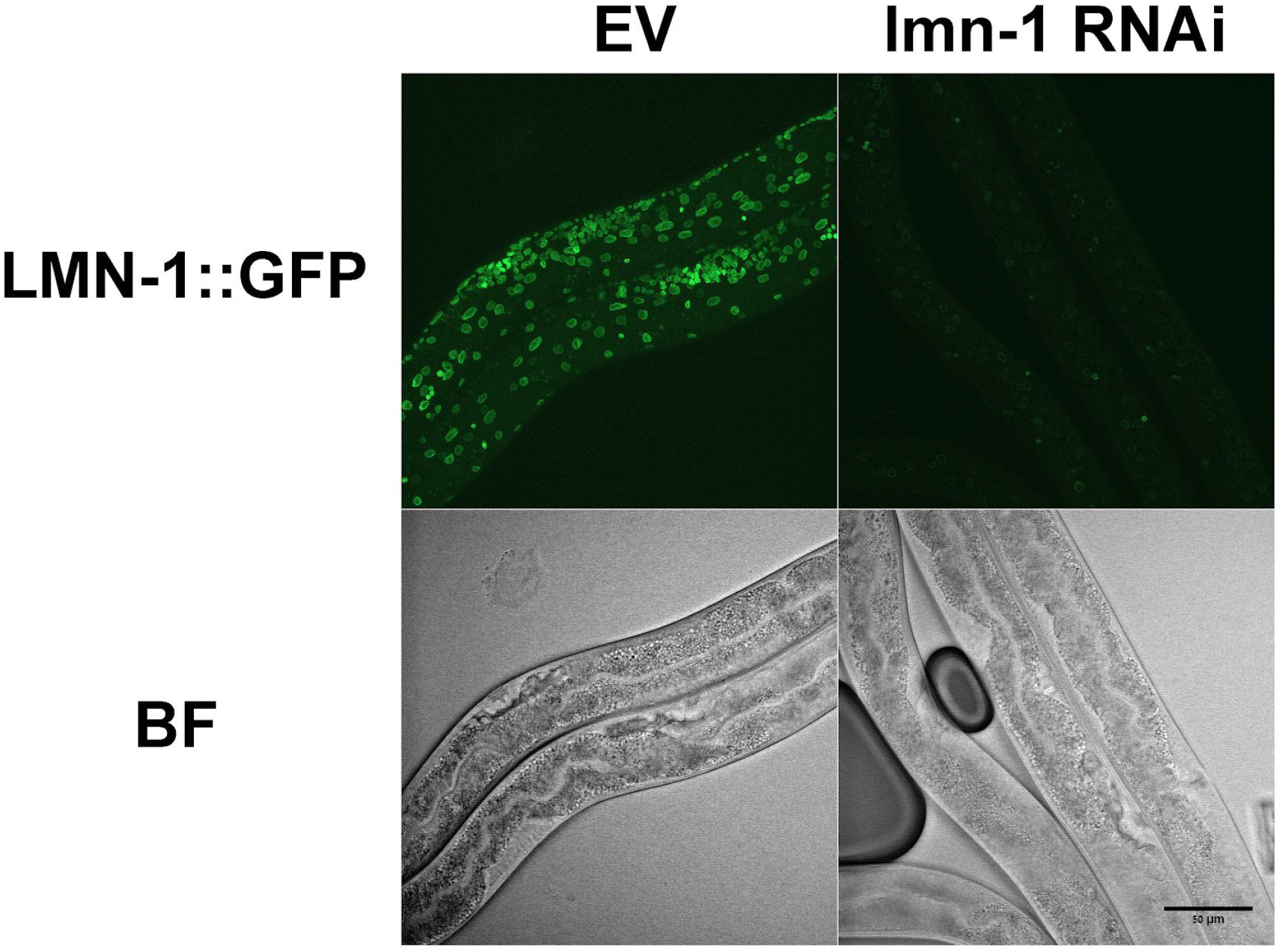
*lmn-1*(RNAi) successfully downregulated lamin, as evident from confocal microscopy of LMN-1::GFP animals.

**Supplementary figure 2.**
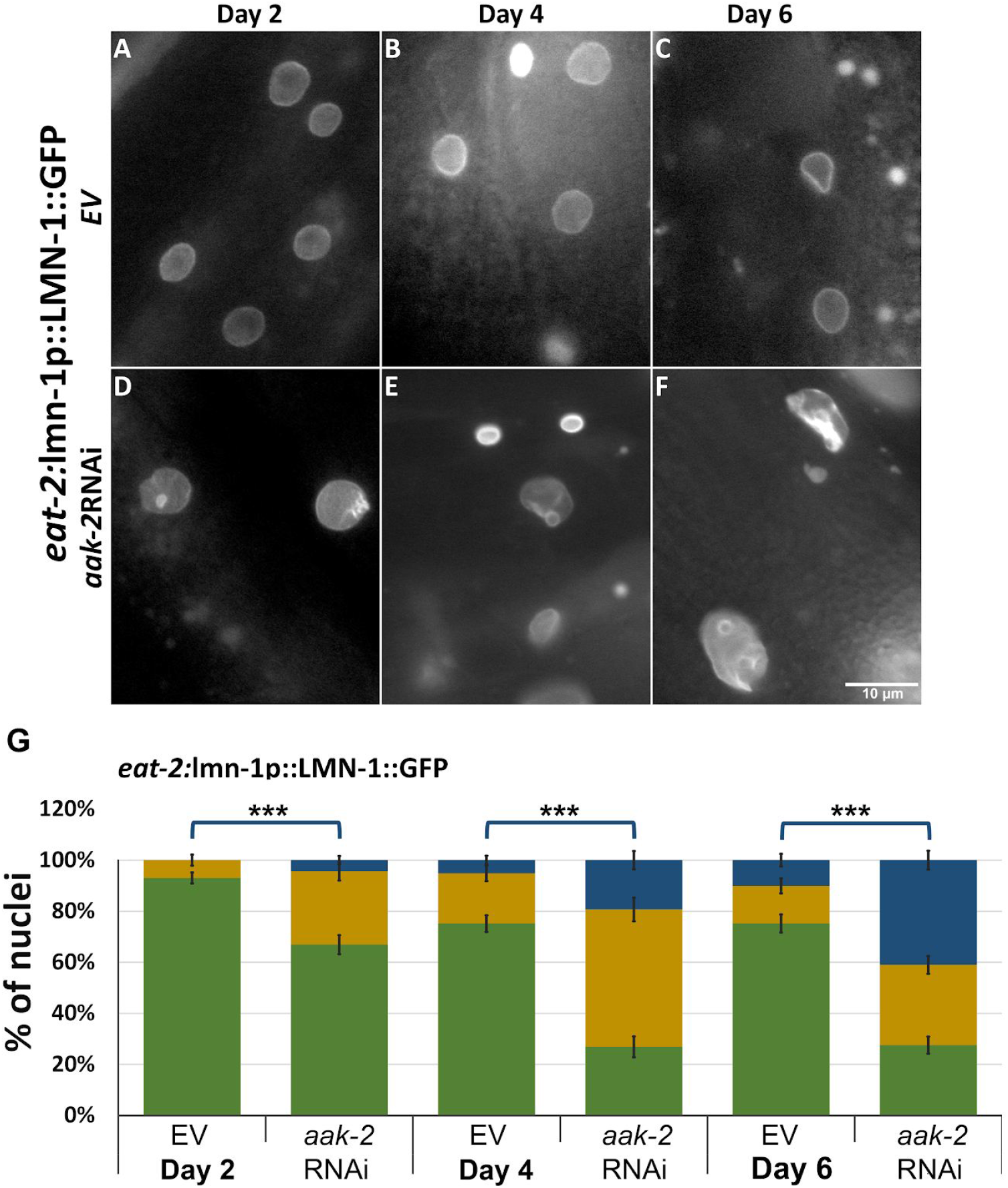
AAK-2 is required to maintain nuclear structure (*A*-*F*) Microscope images of *eat-2* worms expressing LMN-1::GFP fed by either EV (*A*-*C*) or *aak-2* (RNAi) post-hatching (*D*-*F*) at day 2 (*A,D*), 4 (*B,E*) and 6 (*C,F*) of adulthood. (*G*) Relative distribution of the 3 different classes grading the nuclear morphology changes in eat-2 worms expressing LMN-1::GFP fed with *aak-2* (RNAi) or EV at days 2,4 and 6 of adulthood. n(D2)=303 nuclei, n(D4)=292 nuclei, n(D6)=327 nuclei. Error bars in all graphs represent mean ± SEM. P value was calculated using Fisher Exact Probability Test.

**Supplementary figure 3.**
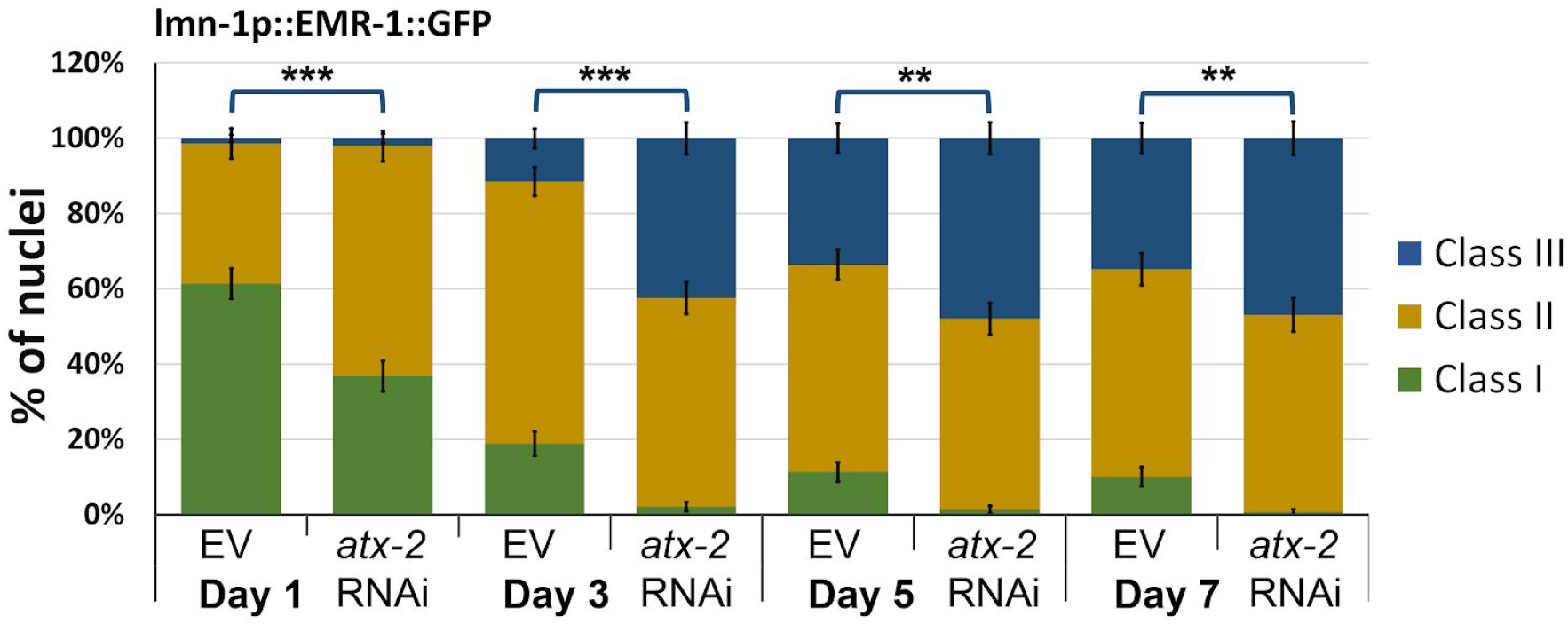
ATX-2 is required to maintain nuclear structure. Relative distribution of the 3 different classes grading the nuclear morphology changes in EMR-1::GFP expressing worms fed with *atx-2* or EV at days 1, 3, 5 and 7 of adulthood. n(D1)=289 nuclei, n(D3)=287 nuclei, n(D5)=291 nuclei, n(D7)=268 nuclei. Error bars in all graphs represent mean ± SEM. P value was calculated using Fisher Exact Probability Test.

